# Identification of Splice Regulators of Fibronectin-EIIIA and EIIIB by Direct Measurement of Exon Usage in a Flow-Cytometry Based CRISPR Screen

**DOI:** 10.1101/2021.05.05.442813

**Authors:** Jessica Hensel, Brent Heineman, Amy Kimble, Evan Jellison, Bo Reese, Patrick Murphy

**Author notes:** Corresponding Author Patrick A. Murphy, Ph.D. Assistant Professor Center for Vascular Biology & Calhoun Cardiology Center University of Connecticut Medical School 263 Farmington Avenue Farmingon, CT 06030.

## Abstract

The extracellular matrix protein fibronectin (FN) is alternatively spliced in a variety of inflammatory conditions, resulting in increased inclusion of alternative exons EIIIA and EIIIB. Inclusion of these exons affects fibril formation, fibrosis, and inflammation. To define upstream regulators of alternative splicing in FN, we have developed an *in vitro* flow-cytometry based assay, using RNA-binding probes to determine alternative exon inclusion level in aortic endothelial cells. This approach allows us to detect exon inclusion in the primary transcripts themselves, rather than in surrogate splicing reporters. We validated this assay in cells with and without FN-EIIIA and –EIIIB expression. In a small-scale CRISPR KO screen of candidate regulatory splice factors, we successfully detected known regulators of EIIIA and EIIIB splicing, and detected several novel regulators. Finally, we show the potential in this approach to broadly interrogate upstream signaling pathways in aortic endothelial cells with a genome-wide CRISPR-KO screen, implicating the TNFalpha and RIG-I-like signaling pathways and genes involved in the regulation of fibrotic responses. Thus, we provide a novel means to screen the regulation of splicing of endogenous transcripts, and predict novel pathways in the regulation of FN-EIIIA inclusion.

## Introduction

Alternative splicing of the essential extracellular matrix protein fibronectin (FN) is tightly regulated during development and in disease ^[1]^. Increased inclusion of alternative exons EIIIA(EDA) and EIIIB(EDB) in the vascular ECM and the circulation coincides with a range of human diseases involving endothelial activation and increased risk of thrombosis and fibrosis, including acute traumatic injury ^[2, 3]^ and chronic responses – as in type II diabetes ^[4, 5]^. Within the vasculature, and in sites of fibrosis, the endothelium is a key source of FN and the FN splice isoforms ^[6, 7]^. Germline deletion of alternative spliced exons EIIIA(EDA) and EIIIB(EDB) leads to lethal vascular defects in development on some genetic backgrounds ^[8]^, and surviving mice exhibit impaired vascular remodeling and risk of vascular hemorrhage ^[8-12]^. Genetic deletion of the EIIIA exon alone reduces clot formation ^[13, 14]^ and fibrotic responses in the skin ^[6, 15]^, liver ^[16]^, lung, ^[17]^ and in heart attack ^[18]^ and stroke models ^[14]^. Thus, EIIIA and EIIIB inclusion is tightly regulated, and is associated with vascular remodelling, thrombosis and fibrosis.

We showed that EIIIA and EIIIB inclusion in the endothelium increases in response to hematopoietic cell recruitment, but the molecular mediators are not yet clear ^[12, 19]^. TGFb is an important mediator of endothelial EIIIA inclusion in fibrotic responses in the liver ^[7, 20]^; however, there are likely other molecular mediators as well. In other cell types, extracellular cues regulating EIIIA inclusion include basement membrane proteins ^[21]^, and growth factors such as HGF ^[22]^. PI3K signaling is an important intracellular mediator EIIIA and EIIIB induction in some contexts ^[23-25]^, leading to increased activity of splice factor phosphorylation and activation. Serine and arginine rich splicing factors (e.g. SRSF1, −3, −5, and −7) promote EIIIA and EIIIB inclusion in an apparently additive manner ^[26-28]^. In contrast to the redundant activities SRSF proteins on EIIIA and EIIIB inclusion, the splice factor Rbfox2 is essential for EIIIB inclusion in multiple cell types ^[19, 29]^, and also has an important role in EIIIA induction in the arterial endothelium ^[19]^. Nevertheless, knowledge of upstream mediators of EIIIA and EIIIB inclusion is still incomplete.

Here, we describe an approach to assess RNA splicing in native transcripts with single cell resolution, and show how this approach allows for the screening of upstream mediators of EIIIA and EIIIB inclusion in cultured aortic endothelial cells. We validate that this approach identifies known regulators of EIIIA and EIIIB splicing in a targeted screen, and use the approach in a genome-wide screen to identify novel genetic dependencies.

## Results

### Direct measurement of FN splicing variants using fluorescent *in situ* probes

We established an assay to examine endogenous RNA splice isoforms in single cells using a commercial *in situ* hybridization protocol compatible with flow-cytometry (PrimeFlow). DNA oligomer probes were designed to bind the alternative fibronectin exons EIIIA and EIIIB, as well exons in the constitutive region of Fibronectin. These probes allow sequence specific fluorescent detection of RNA by flow cytometry after signal amplification (Figure 1A). To validate our approach, we used murine aortic endothelial cells with and without genetic deletion of the EIIIA and EIIIB exons. We found increased staining with both the EIIIA and EIIIB probes in FnAB-het cells, compared to FnAB-null cells (Figure 1B and data not shown), indicating that our probes are specific to these domains. Importantly, there was no difference in total FN staining in the cells, showing that the reduced signal is specific to the splice domains within the FN transcript (Figure 1B). Notably, our results indicate very little background signal from the EIIIA and EIIIB probes in the absence of RNA substrate. The signal for both probes in FnAB-null cells is comparable to cells without the EIIIA or EIIIB probe (Figure 1B). As we were most interested in the ratio of splice isoform to total FN, or the percent spliced in (Psi), we normalized EIIIA and EIIIB inclusion to total FN. We observed that there is a range of inclusion frequencies between cells, but that almost all FnAB-het cells have inclusion levels above the no inclusion controls (FnAB-null, Figure 1C&D). These results confirmed our methodology and provided the basis for CRISPR screening and identification of EIIIA and EIIIB regulators.

**Figure 1.**
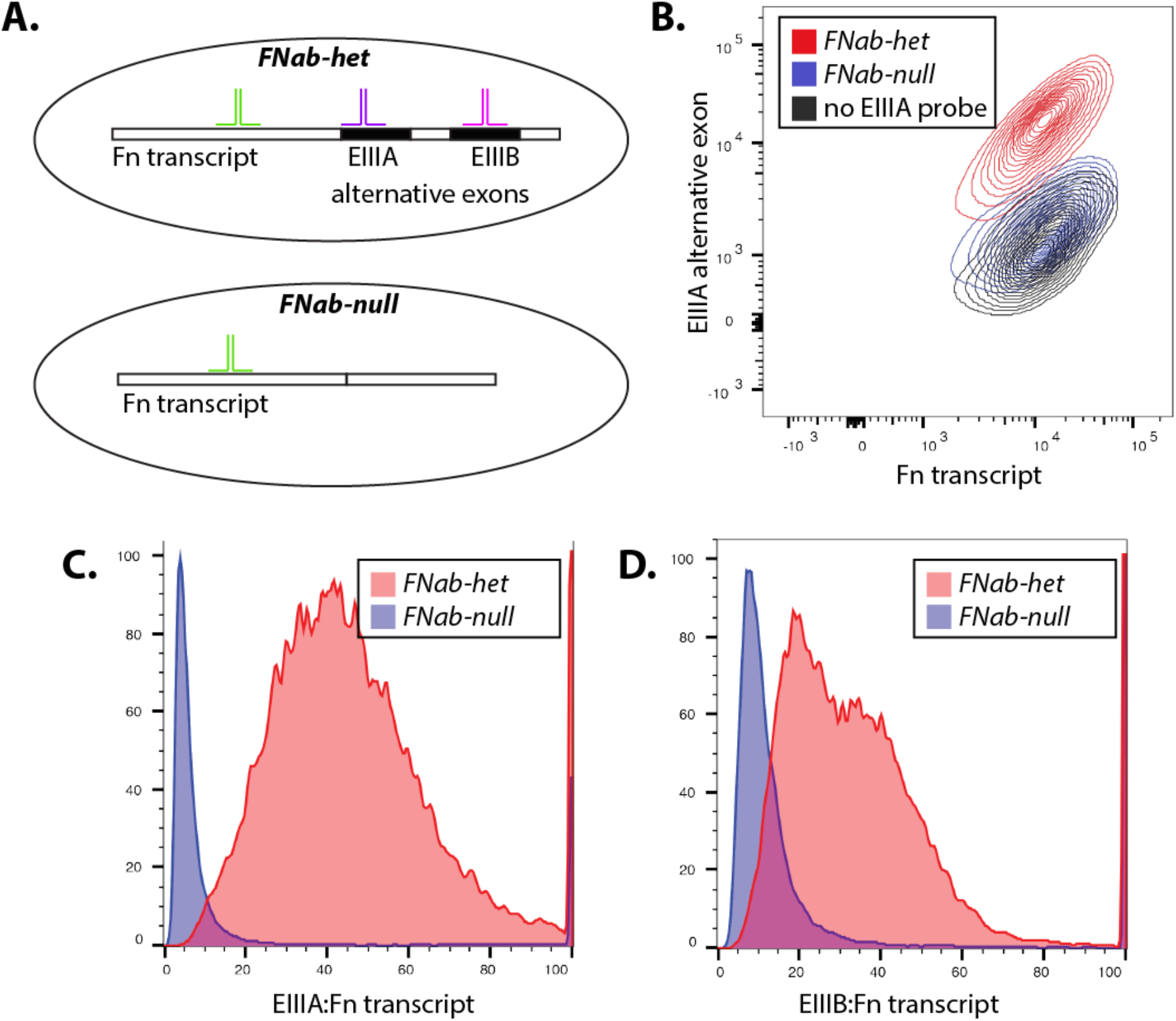
RNA-hybridization assay to measure inclusion of fibronectin alternative exons in individual endothelial cells. (A) Schematic showing binding of probes to transcripts containing EIIIA (from FNab-het cells) or not (from FNab-null cells). Probes to the constitutive region of FN bind to both transcripts (green), but probes to EIIIA (purple) or EIIIB (pink) binds only to these alternatively spliced exon (black region of FN transcript) when they are included in cells. (B) Fluorescent flow cytometry plot showing absorbance detected in all cells for the FN probe (x-axis), but only detected for the EIIIA probe (y-axis) in cells expressing FN with EIIIA (red). Signal is absent when no EIIIA probe is included (but fluorescent detection is still performed, black), or when cells do not produce the EIIIA inclusive FN transcript (blue). (C&D) Ratio of signal from EIIIA probe (A) or EIIIB probe (B) to total FN probe from flow cytometry as in (B).

### Small scale screen of candidate RBPs identifies known regulators of EIIIA and EIIIB

We had observed that EIIIA and EIIIB are co-regulated with hundreds of other skipped exons in endothelial cells activated by hematopoietic cell recruitment under low and disturbed flow ^[12, 19]^. Hypothesizing that common upstream RNA-binding proteins (RBPs) regulate this splicing response, we had assessed flanking regions of these skipped exons to identify likely regulatory RBPs ^[19]^. This identified Rbfox2, which we previously found is required for EIIIB inclusion and contributes to EIIIA inclusion in the endothelium ^[19]^. However, we also identified 57 other RBPs likely active in this response (SI Figure 1). To determine whether any of these RBPs also regulate EIIIA and EIIIB inclusion we performed a small scale CRISPR KO screen, using our *in situ* hybridization protocol as a readout (Figure 2A and SI Figure 2). Five guide RNA (gRNA) pairs were designed to each of the candidate RBPs, and constructs were generated in a pooled approach. Analysis of the plasmid pool showed good representation of each targeting construct and minimal skew across the library (SI Figure 3A&B). Lentivirus was produced in a pooled approach, and transduced into murine aortic endothelial cells at MOI<0.3. A collection of cells at 3 days post-post infection (dpi) showed that representation in cells mirrored the representation in the plasmid library (Figure 3B).

**Figure 2.**
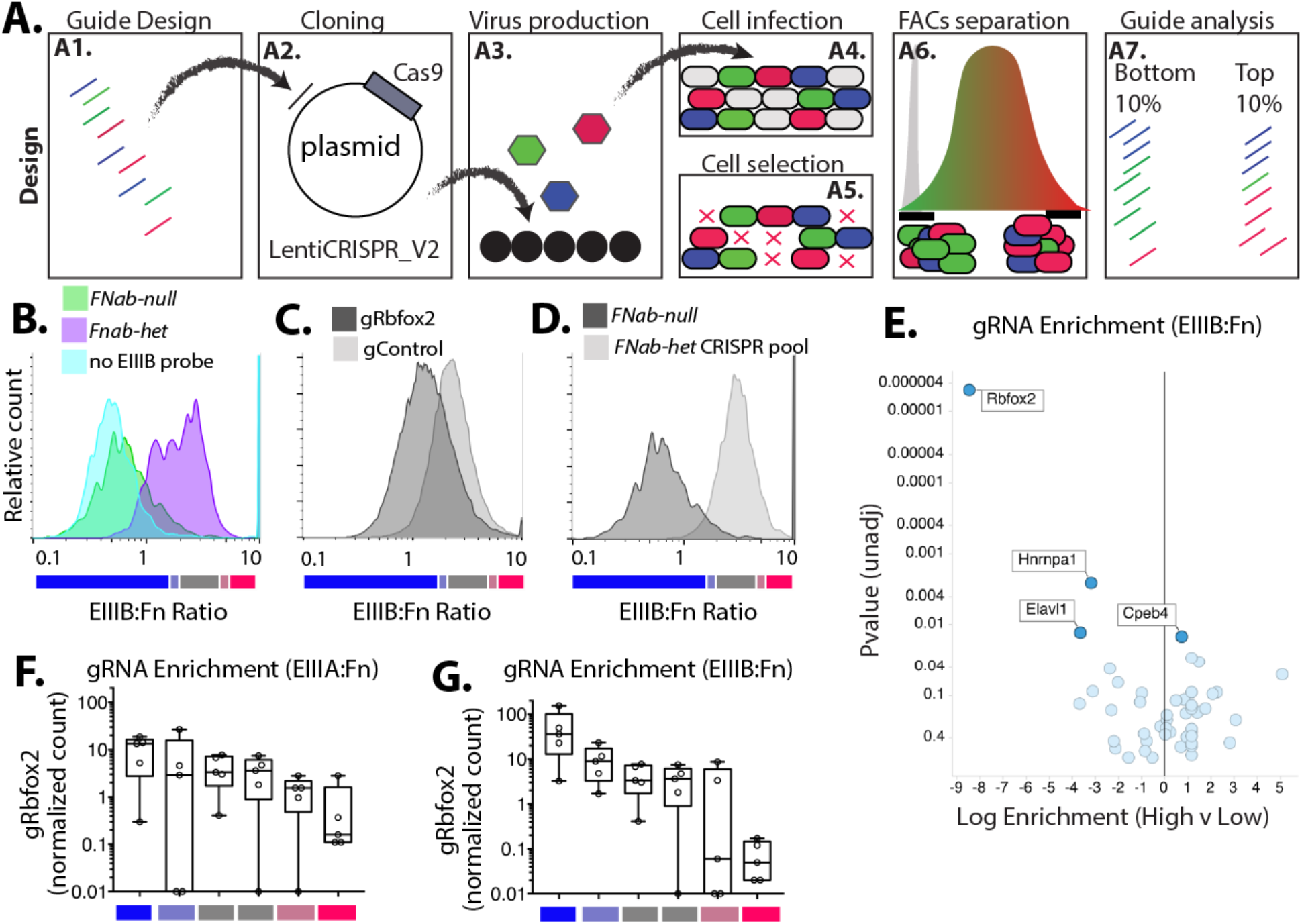
Application of CRISPR screening approach to define regulators of EIIIIA and EIIIB inclusion. (A) Screening approach (A1) Guides to each target gene of interest were designed using the latest rules in the Broad design tool, and prepared as an ssDNA oligo pool. Low-cycle PCR generates dsDNA with ligation sites for (A2) batch cloning into a lentiviral backbone containing Cas9 and guide expression site. (A3) The resultant plasmid library is then used for production of lentivirus and infection of cells (A4). Infection is at 0.3 MOI and (A5) selection is by puromycin. (A6) Selected cells are sorted on the inclusion of alternative exon. Cells in the top 10% and bottom 10% of the response are isolated. (A7) Genomic DNA is isolated and guide enrichment in cells is assessed by sequencing of amplified lentiviral insertions. (B-D) Ratio of EIIIB to Fn probes, showing cells with and without EIIIB or no EIIIB probe at all (B), a guide targeting Rbfox2 (gRbfox2) or a control guide (gControl) (C) and the pool of CRISPR guides targeting all 57 splice factors (D). (E) Volcano plot showing the enrichment of guides for the indicated genes among cells with high levels of EIIIB inclusion versus low levels of EIIIB inclusion. Log enrichment and p-value is from MaGeCK analysis. (F&G) Plots showing the normalized count for individual guides targeted Rbfox2 in each sorted group of cells. Low, mid, or high levels of EIIIA and EIIIB inclusion are shown and represent the bins shown in B-D

**Figure 3.**
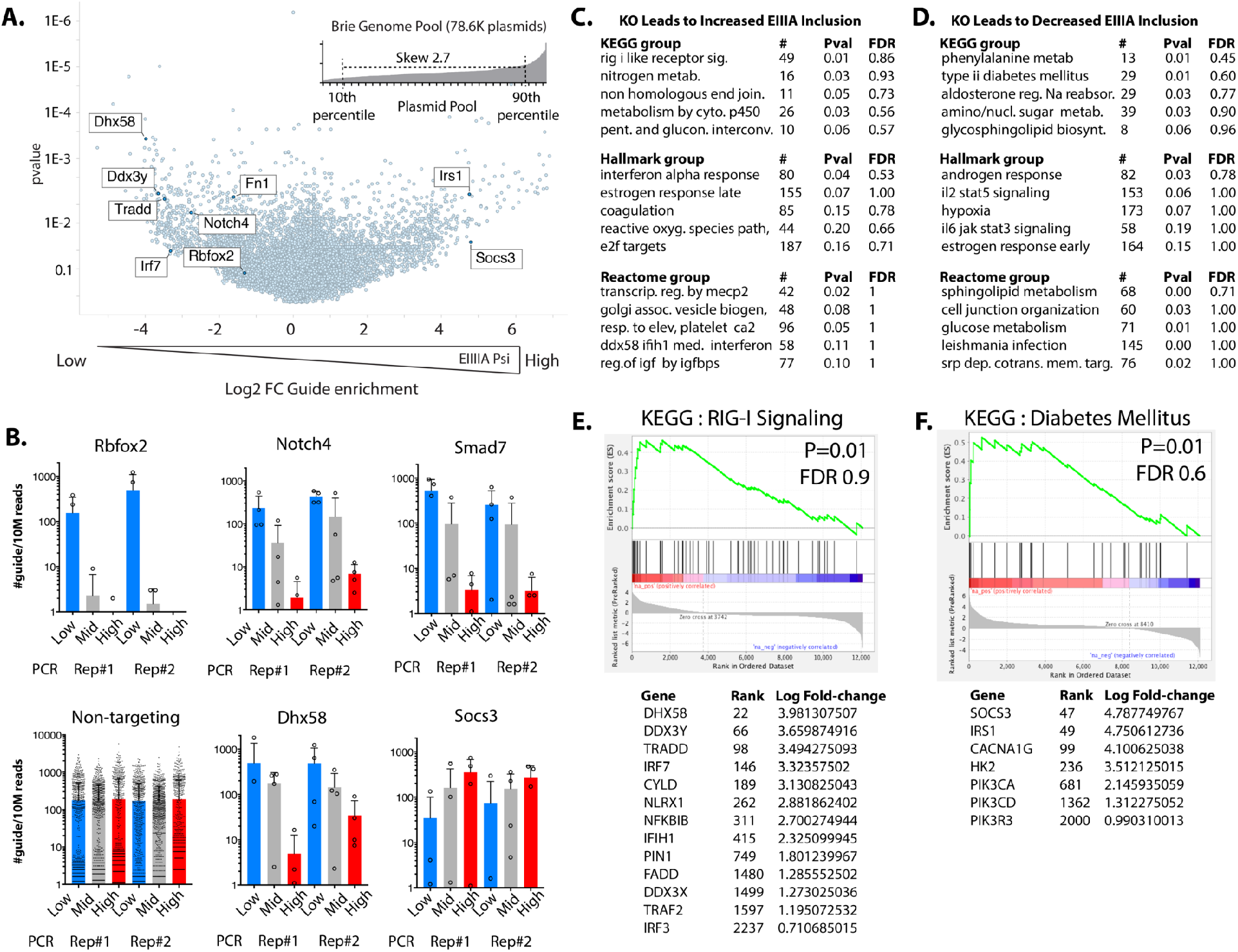
Genome-wide identification of EIIIA regulators. (A) Guide representation in plasmid pool (Brie library) after amplification in bacteria (inset), and volcano plot from MaGeCK analysis showing genes predicted to positively or negatively regulate EIIIA inclusion, based on the enrichment of their specific gRNA. (B) Plots showing individual guides for the genes of interest, and their relative enrichments in sorted cells with low, mid, or high levels of EIIIA inclusion. (C) GSEA analysis showing the enrichment of genes associated with specific KEGG, Hallmark and Reactome gene groups among those targeted in cells with reduced EIIIA inclusion (indicating that these genes promote EIIIA inclusion), and (D) similar analysis of genes targeted more frequently in cells with increased EIIIA inclusion (indicating that these genes suppress EIIIA inclusion). (E) Example of GSEA plot and enriched genes from (C), and (F) similar GSEA plot from (D). GSEA=gene set enrichment. # = number of genes in group. Pval = unadjusted P-value. FDR=False Discovery Rate

Having created a cellular library with constructs targeting each of our candidate RBPs, we then tested for regulation of EIIIA and EIIIB inclusion using our splicing assay. In addition to the FnAB-null cells, we included an efficient lentiviral CRISPR KO construct to Rbfox2 which reduces Rbfox2 protein and EIIIB inclusion as a control (SI Figure 4). This resulted in the expected reduction of EIIIB inclusion, confirming that deletion of a regulatory splice factor leads to a detectable change in splicing (Figure 2). In our CRISPR KO pool, we predicted that Rbfox2 and other gRNA targeting RBPs important in EIIIA or EIIIB inclusion would be enriched in cells with lowest inclusion levels. Conversely, we predicted that gRNA targeting RBPs suppressing EIIIA or EIIIB inclusion would be enriched in cells with highest inclusion. Therefore, we sorted the top 10%, bottom 10%, based on EIIIA or EIIIB inclusion (Figure 2D, and SI Figure 4). We sorted 200K cells in each pool, providing a theoretical coverage of >700 cells per guide (SI Figure 2). Lentiviral insertions were amplified from biological replicates of sorted populations in a nested PCR, as previously reported ^[30, 31]^. Counting and distribution analysis with a negative binomial model was performed using MaGeCK ^[32]^. As expected, we found the strongest signal for Rbfox2 as a required regulator of EIIIB, and a weaker signal as a regulator of EIIIA (Figure 2E-G and SI Figure 5). Also as expected, guides to Srsf5, which promotes of EIIIA inclusion, were enriched in cells with reduced EIIIA (SI Figure 5). We confirmed this effect in cells infected with a single Srsf5 targeting lentivirus (SI Figure 5). Together, these results confirm the detection of expected regulators, and also hint at a role for previously undescribed EIIIA and EIIIB regulators, such as Hnrnpa1 and Elavl1 (EIIIB) and Pcbp2 and Cpsf6 (EIIIA).

### Genome-wide CRISPR KO screen for regulators of EIIIA inclusion

As small-scale screens identified known splice-factor regulators of EIIIA and EIIIB, we aimed to expand this approach to examine the regulation of EIIIA specifically on a genome-wide scale. We used the Brie genome-wide CRISPR-KO library ^[33]^. The plasmid library, which showed good conservation and low skew in the representation of all plasmids, was used to produce virus and infect aortic endothelial cells at >100 cells per guide (at <0.3 MOI), resulting in a cellular library with a skew of 30 from the top 20% to the bottom 80%, and guide coverage of >97%. The library was kept at >100 cells per guide coverage, and two separated pools of virus production and infection were used to generate the final pool (see SI Figure 2). Cells were expanded over two weeks in culture, and were assessed by splicing analysis. The top 10%, bottom 10% and mid 30% of cells were isolated, based on their expression of EIIIA to total FN. Each of these populations was separated into multiple PCR reactions (see SI Table 1 and SI Figure 2), and an Illumina library was generated by nested PCR reactions.

We found that expected genes were enriched. For example, gRNAs to Rbfox2 are enriched among cells with lowest EIIIA inclusion (Figure 3A&B). This is consistent with both the results of our small-scale screen and a reduction in EIIIA with endothelial deletion of the splice factor *in vitro* and *in vivo* ^[19]^ (SI Figure 7). Guides to Notch4 were also enriched among cells with the lowest levels of EIIIA inclusion (Figure 3B), consistent with prior work showing that Notch4 is required for EIIIA expression in lymphatic valves *in vivo*, and in lymphatic endothelial cells *in vitro* ^[34]^. Guides to TGF-beta receptors were not enriched in endothelial cells with low EIIIA inclusion, suggesting that in contrast to other settings, EIIIA responses in aortic endothelial cells may not require sustained TGF-beta signaling. This is supported by our own *in vitro* and *in vivo* data showing no effect of TGF-beta inhibitors on EIIIA inclusion in these aortic endothelial cells (SI Figure 8).

To better understand the pathways regulating EIIIA inclusion, we identified genes promoting EIIIA inclusion or EIIIA skipping based on gRNA enrichment and performed a gene set analysis of enriched genes using GSEA (Figure 3C-F). We found that the deleted genes linked to reduced EIIIA inclusion are concentrated in the RIG-I-Iike signaling pathway (KEGG), and interferon alpha response (Hallmark) – two related inflammatory pathways (Figure 3C&E). We also found a strong signature for genes involved in the regulation of transcription by Mecp2 (Reactome) (Figure 3C). Genes that increased EIIIA inclusion when deleted included Type II diabetes (KEGG), Stat3, and Stat5 signaling pathways (Figure 3D&F), all of which are linked to insulin signaling responses. A suppressive effect of insulin signaling pathways on EIIIA inclusion is consistent with work showing increased EIIIA expression and fibrotic responses in type II diabetes ^[4, 5]^.

### Genes regulating EIIIA inclusion correlate with published fibrotic phenotypes

As EIIIA inclusion is a driver of fibrotic responses, we predicted that deletion of genes which suppress EIIIA inclusion would lead to reduced fibrosis. Conversely, we expected that deletion of genes which lead to increased EIIIA inclusion would increase fibrosis. To test this, we asked whether EIIIA-regulating genes also regulate fibrotic responses in published data (Figure 4A-B). We examined the top 100 gene deletions predicted to increase EIIIA inclusion (“High EIIIA”) or promote EIIIA skipping (“Low EIIIA”) based on the enrichment of their gRNA and an unadjusted p-value <0.05. We performed a literature search for effects of gene deletion or suppression on fibrotic phenotypes *in vivo* or *in vitro*. We found literature support for the effects of KO or KD on fibrosis (increased or decreased) for 14% of the genes in both Group I and Group II. CRISPR-targeted genes leading to increased EIIIA inclusion (High EIIIA) were associated with increased fibrosis in KO or KD (84% increased fibrosis, Figure 4A). The CRISPR-targeted genes which led to reduced EIIIA inclusion in our screen (Low EIIIA), KO, or KD are associated mainly with less fibrosis (71% reduced fibrosis, Figure 4A). While we assessed only a few of the genes in our screen, we would predict that many of the other still unannotated genes will have similar effects on the fibrotic response (Figure 4B). Thus, our data predicts genes involved in the fibrotic response *in vivo* and *in vitro*.

**Figure 4.**
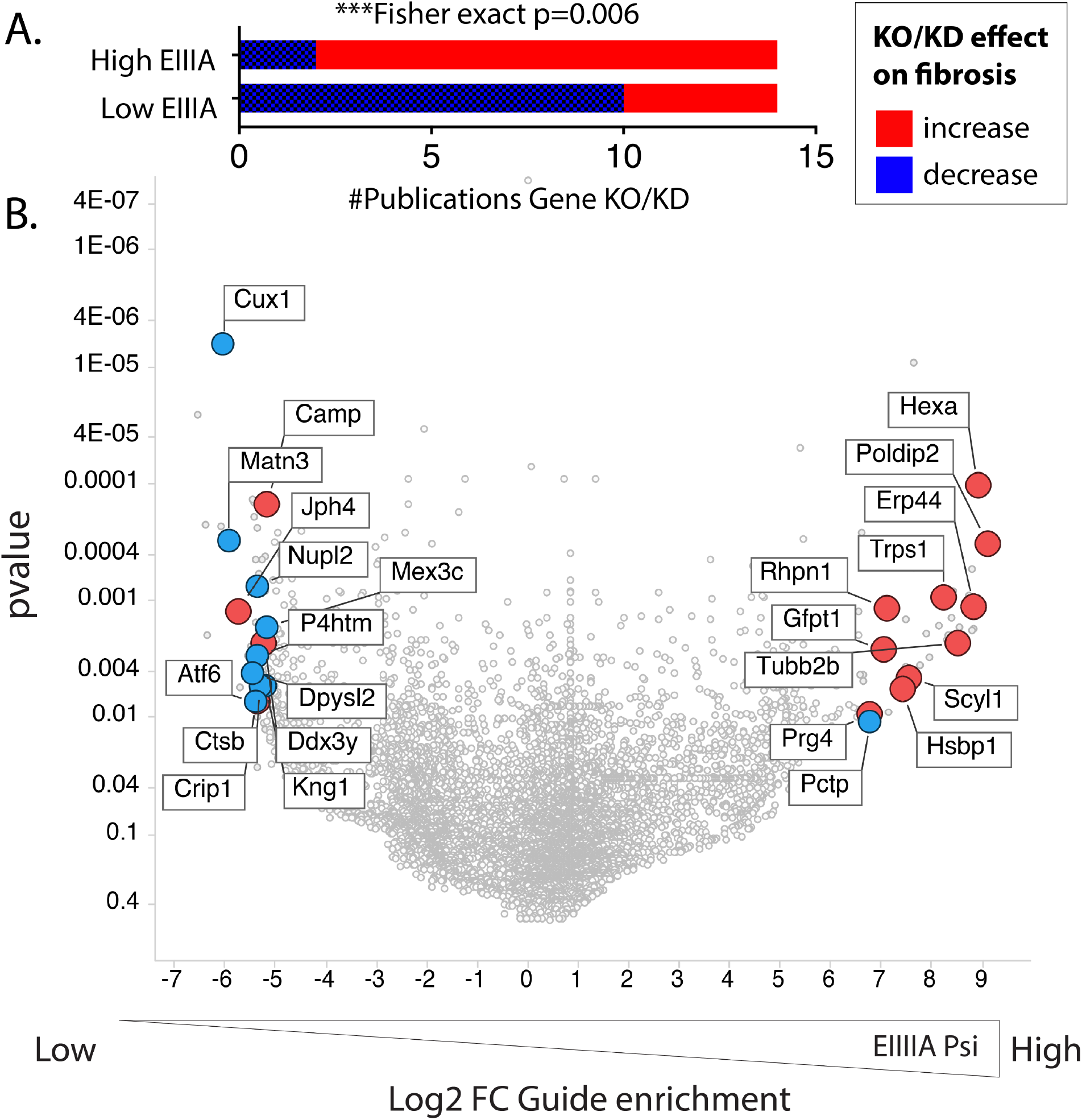
FN splicing regulators correlate with fibrotic phenotypes. (A-B) Analysis of top regulatory genes from CRISPR screen in publicly available data. (A) Graph showing the literature supporting either an increase or decrease in fibrotic effects in mice or cells in culture, following gene deletion or suppression. The top 100 most enriched genes in cells with either increased EIIIA inclusion or decreased EIIIA inclusion were assessed. (B) Volcano plot, showing the genes with literature support for effects on fibrotic responses, positively or negatively

## Discussion

Here, we validate a novel means of assessing splicing responses in individual cells by multicolor *in situ* hybridization and show the utility of this approach in a CRISPR KO screen for upstream regulators of EIIIA and EIIIB inclusion. This approach addresses a number of limitations with splice reporters commonly used for this type of screen by preserving contextual and chromosomal specific regulatory mechanisms closely tied to the spliceosomal machinery, and allowing the analysis of primary cells without the introduction of retroviral or plasmid splice reporter constructs. Furthermore, we harness this approach for the broad and unbiased identification of novel regulators of FN-EIIIA splicing, identifying a strong positive correlation with genes affecting fibrotic response in published data.

### Advantages and limitations of multicolor *in situ* hybridization over splice reporters

Our approach is the first to assess splicing changes within individual cells in a flow-cytometry based assay. To identify regulators of alternative splicing, prior work used splice reporters as a surrogate for the endogenous splicing change. These reporters typically include the alternative exon and some portion of the upstream and downstream introns and adjacent exons, and lead to the expression of a downstream fluorescent or luciferase protein reporter when the exon is spliced in ^[35-38]^. While these reporters allow the testing of intronic and exonic sequences contributing to splicing changes, they suffer from a range of disadvantages that limit their accuracy as surrogates for endogenous splicing patterns. First, as the reporters are typically plasmid or retroviral insertions, they are size restricted, and lack the chromosomal context of the actual gene location. There are extensive interactions between gene promoters, transcriptional machinery, and RBPs regulating the spliceosome ^[39]^. There are also direct impacts of transcriptional regulation on RNA splicing ^[26]^. These factors limit a splicing reporter’s ability to accurately recapitulate the contextual machinery that we know contributes to the regulation of alternative splicing. Second, RNA splicing can be regulated by long-range interactions within the mRNA itself, sometimes several kb from the regulated splicing event ^[40]^. Given the size limitations of splicing reporters introduced into cells, these long-range interactions are not typically considered. Third, as splice reporters must be introduced into cells, it is difficult to work with primary cells and most work has been done with plasmids in easily transfected cell lines, which may not be relevant to the cellular phenotype being modeled. Fourth, as these reporters also measure the stability of the protein readout, effects on protein stability must be carefully controlled for in assay design ^[36]^. Thus, by several measures, our direct *in situ* hybridization screen offers benefits over splice reporter screens.

Although *in situ* hybridization screening has benefits, it also has some limitations. First, this assay is capable of detecting splice isoforms only in fixed and permeabilized cells; thus, iterative screens to purify cells with most or least inclusion is not possible. Second, the splicing change must be large enough to land the oligomer probes on the novel sequence. The EIIIA and EIIIB exons are ~270bp in length, but shorter regions would be much more challenging. Lastly, the sensitivity of the approach depends not only on size of the targeted region, but also the copy number in the cell. This assay is reported to have a sensitivity of below 10 transcripts per cell in optimal conditions ^[41]^, but given the additional size limitations, sensitivity to splice isoforms is likely less. FN is among the most expressed transcripts in our aortic endothelial cells, with expression level similar to beta-actin (by FPKM). EIIIA and EIIIB exons contribute to the majority of these transcripts in the aortic endothelial cells. Therefore, FN and the alternative exons EIIIA and EIIIB provide abundant targets for this approach. Novel methods to detect and quantify RNA transcripts in cells may be needed to adapt this assay for the analysis of lower abundance splice isoforms.

### Identification of novel pathways modulating EIIIA inclusion

This new data further validates our previous discovery that Rbfox2 is a key regulator of EIIIB inclusion in arterial endothelial cells ^[19]^, as it is in mouse stem cells ^[29]^. Here, we also show that Rbfox2 affects EIIIA inclusion, although the effects on EIIIA are not as pronounced as on EIIIB, which is also consistent with our prior observations in Rbfox2 KO mice ^[19]^. Among other known regulators of FN splicing, Srsf1 and Srsf3 are notably absent, possibly due to impaired cellular growth or increased lethality caused by complete deletion. Indeed, we have found that protein expression is retained in cells with Srsf1 targeting guides, suggesting that these guides are either inactive, or more likely, that the indels they generated have retained the Srsf1 protein coding sequence (SI Figure 6). However, Srsf5 deletion appears to be well tolerated, and its deletion suppressed EIIIA inclusion in our screen and also in cells when measured by qPCR (SI Figure 5). Although we have not yet tested other predicted splice factor regulators of EIIIA and EIIIB inclusion, these are new candidates for the regulation of this splicing response.

Genes predicted to regulate EIIIA inclusion by our screen often regulated fibrotic responses in the same direction, so that increased EIIIA inclusion with gene deletion correlated with increased fibrosis in mouse models or cell culture (Figure 4). While this is consistent with the profibrotic effects of EIIIA inclusion in animal models, it is difficult to say whether these effects are due to EIIIA inclusion alone, or more likely, that deletion of these genes promotes a profibrotic response in endothelial cells leading to EIIIA inclusion. For example, Hsbp1 was identified as a negative regulator of EIIIA inclusion in our screen (KO leads to increased inclusion), and a negative regulator of fibrosis and endothelial-mesenchymal transition (KO leads to increased fibrosis and endMT) ^[42]^. In lung fibrosis models, overexpression of Hsbp1 specifically in the endothelium strongly suppressed collagen deposition in the lung ^[42]^. Conversely, Mex3c was identified as a positive regulator of EIIIA in our screen and promotes fibrotic responses. Mex3c promotes PTEN^K27-polyUb^ ubiquitination, which is in turn important for PTEN activity and phosphorylation and accumulation of TWIST1, SNAI1, and YAP1, which ultimately leads to epithelial-mesenchymal transition and fibrosis ^[43, 44]^. As EIIIA inclusion is a known regulator of fibrotic responses, it would be interesting to determine the relative contribution of EIIIA splicing changes in the responses. Furthermore, a focused screen examining the ~85% of other genes we find to regulate EIIIA, but not yet linked to fibrotic responses, may reveal new regulators of fibrosis.

In conclusion, we have demonstrated that the direct detection of splice isoforms can be used to identify regulators of alternative splicing in CRISPR screens. Using this approach, we have performed the first genome-wide screen for regulators of FN-EIIIA, revealing candidate pathways in the regulation of EIIIA inclusion and fibrotic responses in endothelial cells.

## Materials & Methods

### Isolation and immortalization of mouse aortic endothelial cells (mAECs)

Mouse aortic endothelial cells were isolated from FnAB-null and -het mice as described in Murphy et al. 2018 ^[19]^. Cells were then sorted by flow cytometry (FACs) (BD Aria) on acetylated LDL+ and endothelial cell markers Icam2+ and Cd31+. After expansion, endothelial cells were immortalized by lentiviral TetOn-SV40 using the VSVG packaging plasmid. Cells were grown in low glucose DMEM and 10% FBS under 3% O2 conditions using 2ug/mL doxycycline (Dox).

### Generation and amplification of CRISPR plasmid pools

(Splice factor pool) Five CRISPR-KO guide sequences were designed to each of 57 splice factor targets using described rules (Azimuth 2.0) ^[33]^. Oligos were ordered and BsmBI batch cloned into the gRNA expression cassette of a LentiCRISPR_v2 plasmid containing Cas9 (Addgene 52961). The library was amplified in Stbl3 (Thermo) and tested by miSeq analysis for coverage and skew. Individual guides (e.g. to Rbfox2) followed the same protocol, but were confirmed by sanger sequencing. (Genome-wide pool) For the genome wide screen, the Brie library in LentiCRISPR_v2 (Addgene 73632) was obtained as a plasmid library and amplified in Stbl4 (Thermo) on 500cm^2^ LB-agar coated ampicillin+ selection plates. The library was tested by NextSeq analysis for coverage and skew.

### Transduction of Splice Factor pool and Genome-Wide CRISPR pool into mAECs

Lentivirus was generated in 293T cells using delta 8.9^[45]^ and pHCMV-EcoEnv (Addgene 15802)^[46]^ as packaging plasmids and 25kDa linear PEI as a transfection agent. Media was changed at 1 day and supernatant was taken at 3 days after transfection for the treatment of recipient cells at MOI <0.3 (>1000 infected cells per guide). Virus supernatant was added to recipient mAECs with polybrene (8ug/mL), and cells were selected with puromycin for 4 days, confirming a MOI<0.3 and complete killing of uninfected cells. An aliquot of cells was taken 3-4 days post infection for an early timepoint analysis of lentiviral library representation.

### PrimeFlow Detection of Alternative Splicing Isoforms

mAECs harboring CRISPR guides were trypsinized and split into low glucose DMEM and 10% FBS into new dishes, without doxycycline to stop growth. 24 hours later, cells were once again trypsinized and collected for PrimeFlow. Protocol for RNA detection was followed, according to manufacturer’s instructions (ThermoFisher). Protocol was completed over the course of 2 days, using the stopping points described. Target probes were Alexa Fluor 488 for Fibronectin and Alexa Fluor 647 for either EIIIA or EIIIB. FnAB-null cells were included as a control for staining.

### Enrichment of guides in cells with increased or decreased EIIIA or EIIIB inclusion

mAECs with PrimeFlow staining were sorted based on the ratio of EIIIA or EIIIB to total FN, as a measure of inclusion frequency. Samples were digested overnight with Proteinase K at 55C, and DNA collected by Quick DNA kit (Zymo). For the genome-wide screen, at least 3 replicates were taken from each condition (see SI table 1), and isolated independently. Lentiviral insertions in genomic DNA were amplified by nested PCR, using PCR1 primers (see table) followed by PCR2 primers containing barcodes and Illumina priming sequences (see table). PCR reactions were separated using a 2% agarose gel, and bands of interest were excised. PCR products were column cleaned (Gel cleanup kit, Zymo), and pooled based on relative concentration. Pooled samples were sequenced on Illumina NextSeq instrument using 75bp single end (SE) reads.

### Identification of top Fn-EIIIA regulators (Bioinformatics approach)

Fastq files were analyzed using Mageck-0.5.6 software^[32]^, providing counts of each guide in each sequenced population, and differences in their representation (mageck test).

### RNA Extraction and qPCR

RNA extraction was performed using the Qiagen RNAeasy minicolumns, following the protocol provided by the manufacturer (including DNase treatment), and eluting in 10uL of DNase, RNase free water. RNA concentrations were spot-checked (ThermoFisher, Nanodrop 2000) to ensure the RNA extraction was successful. cDNA was prepared using random primers and qPCR was run using BioRad’s iQ SYBR Green Supermix.

Key Resources Table.

Key Resources Table
| Reagent Type | Description | Source |
| --- | --- | --- |
| Vector | LentiCRISPR_v2 | Addgene |
| Packaging Plasmid | Delta 8.9 |  |
| Packaging Plasmid | CMV-Eco | Addgene |
| CRISPR pool | Mouse Genome-wide<br>CRISPR pool | Addgene |
| Commercial kit | PrimeFlow RNA<br>Detection Kit | ThermoFisher |
| Commercial kit | Quick DNA Kit | Zymo |
| Commercial kit | Gel Cleanup Kit | Zymo |
| Software, algorithm | Mageck-0.5.6 | SourceForge |
| Bacterial cells | Stbl3 (targeted screen)<br>Stbl4 (Brie genome-wide screen) | ThermoFisher |
| Commercial kit | RNAeasy minicolumns | Qiagen |
| qPCR Reagent | iQ SYBR Green<br>Supermix | BioRad |

## Supporting information

CountTables

## Acknowledgments

We are grateful for the help of Christopher (“Kit”) Bonin and Geneva Hargis in the UCONN Medical Science Writing group for editing.

## Ethics Approval and Consent to Participate

No animals or human tissues were used in this study.

## Author Contributions

P.M. developed the study concept and design; P.M. performed development of methodology. P.M. and J.H. wrote, reviewed and revised the paper; P.M., J.H., B.H. and A.K. provided acquisition of data. P.M. provided analysis and interpretation of data, and statistical analysis; P.M., J.H., B.H., A.K., E.J. and B.R. provided technical and material support. All authors read and approved the final paper.

## Data Availability Statement

Data from CRISPR screens are included as supplemental information.

