## Supplemental Figures for "Identification of Splice Regulators of Fibronectin-EIIIA and EIIIB by Direct Measurement of Exon Usage in a Flow-Cytometry Based CRISPR Screen"

A.

Induced inclusion of alternative exon with innate immune recruitment

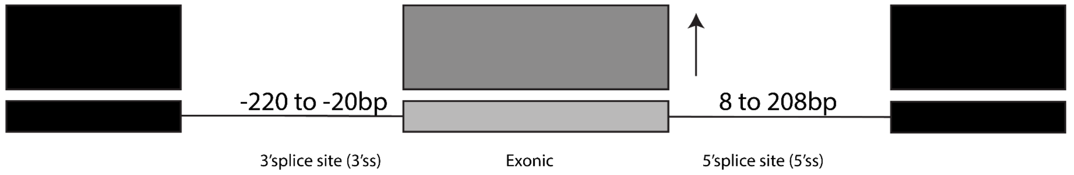


| Induced-3'ss | FDR q-val | FWER p-val |
|---|---|---|
| RBM4B ENSMUSG00000056951 | 0.001186047 | 0.001 |
| CPEB4 CPEB3 | 5.93E-04 | 0.001 |
| U2AF2 | 0.001177889 | 0.003 |
| SRSF7 (human) | 8.83E-04 | 0.003 |
| CPEB2 | 7.07E-04 | 0.003 |
| PPRC1 | 7.82E-04 | 0.004 |
| ANKHD1 ANKRD17 ENSMUSG00000089 | 0.005609915 | 0.035 |
| CNOT4 | 0.010228904 | 0.07 |
| RALY | 0.01151817 | 0.087 |
| ZCRB1 | 0.010604077 | 0.089 |
| PTBP1 PTBP2 ROD1 | 0.009765377 | 0.09 |
| RBM8A B020018G12RIK | 0.008951596 | 0.09 |
| RALYL HNRNPC | 0.00852869 | 0.093 |
| ZC3H10 | 0.008156597 | 0.096 |
| ELAVL1 | 0.007690097 | 0.097 |

| Induced-exonic | FDR q-val | FWER p-val |
|---|---|---|
| RBM4B ENSMUSG00000056951 | 8.33E-04 | 0.001 |
| DAZAP1 | 4.17E-04 | 0.001 |
| PPRC1 | 2.78E-04 | 0.001 |
| SRSF7 (human) | 2.08E-04 | 0.001 |
| SF3B4 | 0.001634647 | 0.01 |
| SRSF2 | 0.001472269 | 0.011 |
| RBMS3 | 0.001261945 | 0.011 |
| ENSMUSG00000021927 HNRNPA1 HNRNPA3 GM8 | 0.001600816 | 0.016 |
| RBM8A B020018G12RIK | 0.00192553 | 0.022 |
| SRSF10 SRSF12 | 0.004452667 | 0.055 |

| Induced-5'ss | FDR q-val | FWER p-val |
|---|---|---|
| CELF3 | 0 | 0 |
| EIF2S1 | 0 | 0 |
| CELF4 | 0 | 0 |
| RBFOX1 RBFOX3 RBFOX2 | 0 | 0 |
| ZCRB1 | 0 | 0 |
| CELF6 | 3.46E-04 | 0.002 |
| RBM4B ENSMUSG0000005695 | 2.97E-04 | 0.002 |
| HNRNPK GM7964 | 0.001803027 | 0.014 |
| SRSF7 (human) | 0.001836342 | 0.016 |
| ELAVL1 | 0.001871341 | 0.018 |
| PCBP1 | 0.001980323 | 0.021 |
| ANKHD1 ANKRD17 ENSMUSG( | 0.002096575 | 0.024 |
| U2AF2 | 0.003849339 | 0.046 |
| PCBP3 PCBP4 | 0.005633013 | 0.071 |
| PPRC1 | 0.006057801 | 0.082 |

B.

Suppressed inclusion of alternative exon with innate immune recruitment

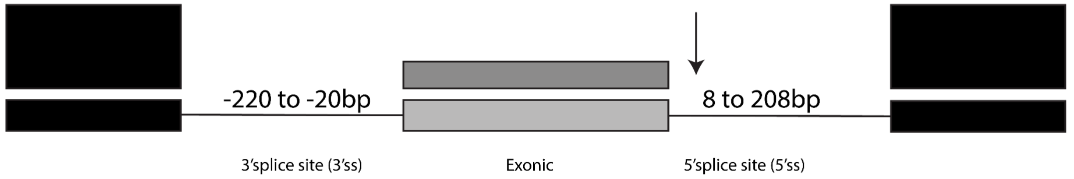


| Suppressed-3'ss | FDR q-val | FWER p-val |
|---|---|---|
| U2AF2 | 0 | 0 |
| RALY | 0 | 0 |
| ELAVL1 | 0 | 0 |
| CPEB4 CPEB3 | 0 | 0 |
| TIAL1 TIA1 | 0 | 0 |
| RALYL HNRNPC | 0 | 0 |
| PTBP1 PTBP2 ROD1 | 0 | 0 |
| CELF3 | 0 | 0 |
| RBM4B ENSMUSG00000056951 | 0 | 0 |
| PCBP3 PCBP4 | 1.29E-04 | 0.001 |
| RBM24 | 2.13E-04 | 0.002 |
| PPRC1 | 4.91E-04 | 0.005 |
| CELF4 | 4.53E-04 | 0.005 |
| PCBP1 | 4.21E-04 | 0.005 |
| HUMAN-SRSF7 | 0.002085737 | 0.027 |
| CPEB2 | 0.002856554 | 0.039 |
| RBM38 | 0.003806701 | 0.054 |
| ANKHD1 ANKRD17 ENSMUSG00000089 | 0.004778821 | 0.072 |
| RBM42 | 0.006076762 | 0.095 |

| Suppressed-exonic | FDR q-val | FWER p-val |
|---|---|---|
| PABPN1 | 0 | 0 |
| CNOT4 | 0 | 0 |
| YBX1 CSDA | 0.001006892 | 0.003 |
| SART3 | 0.001494747 | 0.006 |
| PABPC1 PABPC6 PABPC2 | 0.001815933 | 0.009 |
| HNRPLL | 0.005136305 | 0.031 |
| SRSF10 SRSF12 | 0.004815856 | 0.034 |
| HNRNPL | 0.008012875 | 0.062 |
| SRSF1 ENSMUSG00000090049 | 0.011674899 | 0.1 |

| Suppressed-5'ss | FDR q-val | FWER p-val |
|---|---|---|
| CELF4 | 0.001309524 | 0.001 |
| RALY | 0.003265298 | 0.005 |
| TARDBP | 0.002176865 | 0.005 |
| CPEB2 | 0.001945149 | 0.006 |
| CELF3 | 0.001556119 | 0.006 |
| U2AF2 | 0.001296766 | 0.006 |
| RBM24 | 0.001974098 | 0.011 |
| HUMAN-SRSF7 | 0.002744222 | 0.018 |
| PTBP1 PTBP2 ROD1 | 0.002841946 | 0.021 |
| ELAVL1 | 0.003155093 | 0.026 |
| TIAL1 TIA1 | 0.002962607 | 0.027 |
| SART3 | 0.002805592 | 0.028 |
| CELF6 | 0.002973515 | 0.032 |
| RBM45 | 0.003462037 | 0.04 |
| PABPC1 PABPC6 PABPC2 | 0.005322549 | 0.065 |
| CPEB4 CPEB3 | 0.005838832 | 0.076 |
| PABPC5 | 0.006703992 | 0.092 |

**SI Figure 1. Identification of splice factors active in the response of the endothelium to low and disturbed flow.** Motif enrichment analysis for binding sites of splice factors (from CisBP-RNA database) using GSEA, as described in Murphy et al., 2018 eLife. The top 30 6-mer binding sites for each RNAbp were used to determine enrichment of these RNAbp specific motifs among 6-mers enriched in the selected regions in regulated exons versus non-regulated exons. (A) Analysis of regulated skipped exons induced by low flow and macrophage recruitment. (B) Analysis of regulated skipped exons suppressed by low flow and macrophage recruitment. Top ranking RNAbp from this list were combined with differentially expressed RNAbp and additional known regulators of EIIIA and EIIIB (e.g. Srsf5) to generate the list of target RNAbp for the small-scale screen.

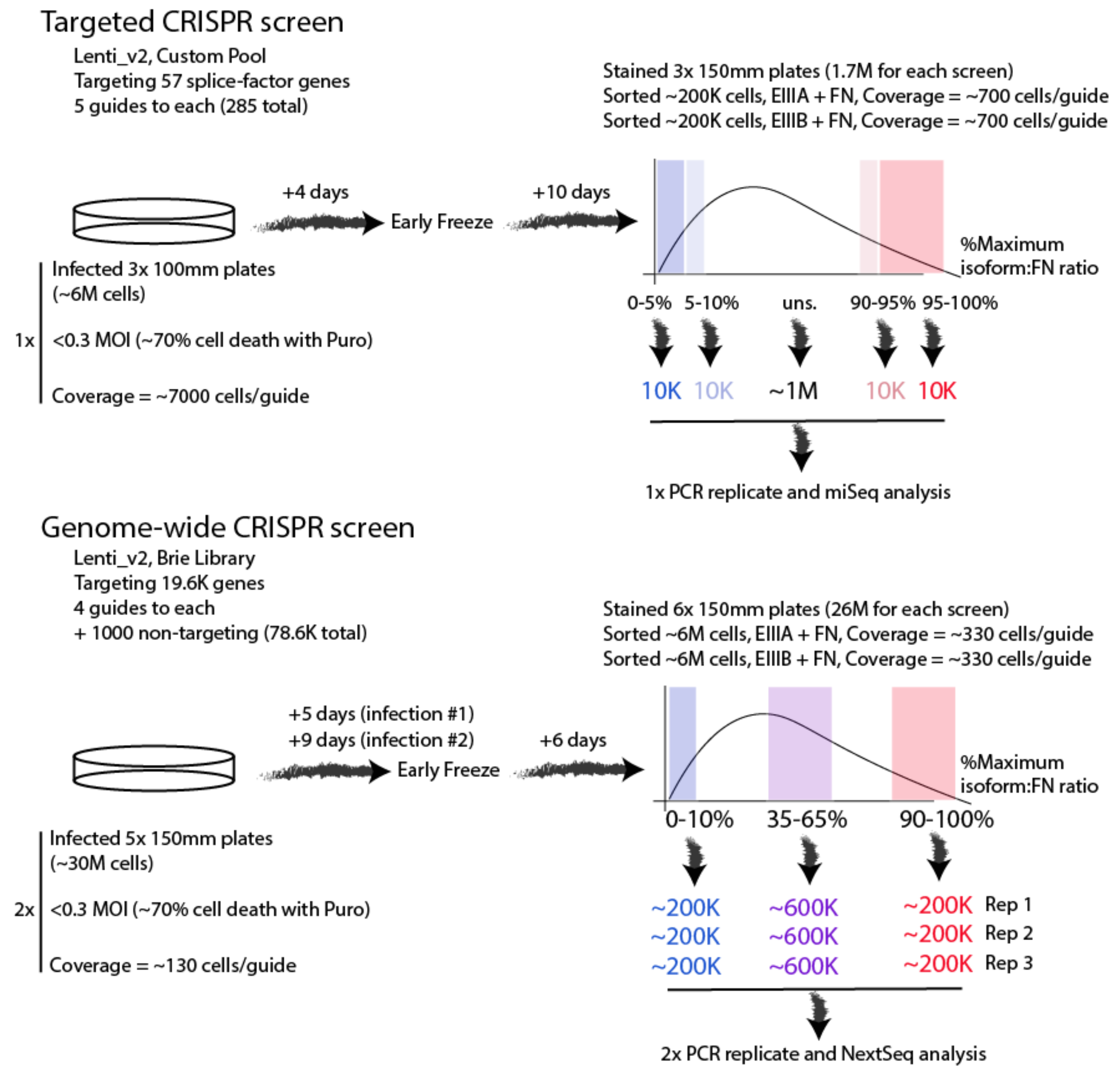


**SI Figure 2. Outline of CRISPR screening approaches.** For both the targeted screen and the larger scale genome-wide screen, cells were infected with virus derived from lentivirus expressing the indicated libraries at <0.3 MOI and selected under puromycin for stable integration of lentiviral constructs. Cell cultures were kept at sufficient size to maintain a coverage of ~7000 cells per guide for the targeted screen and ~100 cells per guide for the genome-wide screen. For the genome-wide screen, this was done in two separate infections, which were then pooled. Within ~2 weeks of infection, cells were stained for EIIIA and FN, or EIIIB and FN, according to the PrimeFlow protocol. The indicated bins were sorted, based on the ratio of the isoform specific (EIIIA or EIIIB) probes to the total FN probe. For the genome-wide screen, several replicates of each bin were obtained, for biological replicates. For the genome-wide screen, the PCR of insertions was performed in two separate PCR replicates, to control for PCR error.

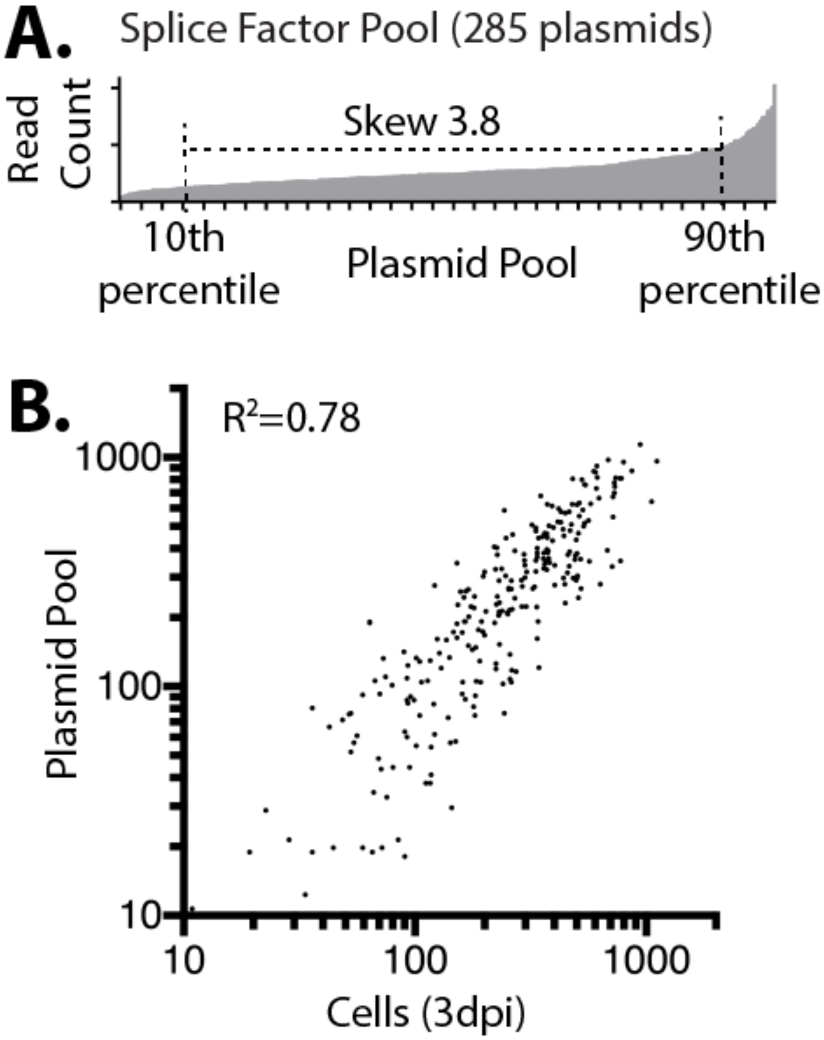


**SI Figure 3. Targeted pool representation in cells.** (A) Read density (y-axis) for each guide construct in the plasmid library for a list of 57 candidate splice factors (5 guides for each gene, x-axis). (B) Read count for the plasmid pool (A, y axis) and cells after infection with lentivirus derived from the plasmid pool (x axis). Each point represents a single gRNA sequence.

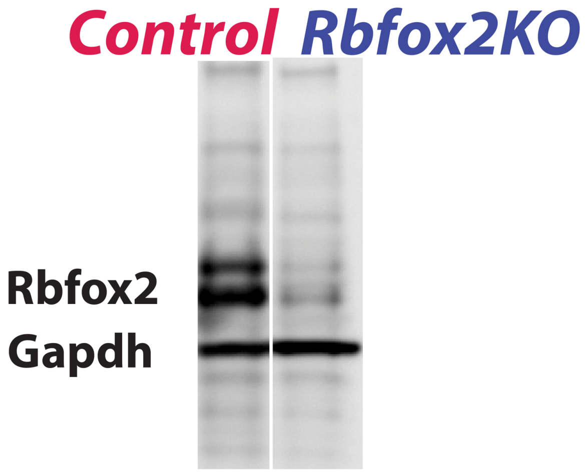


**SI Figure 4. CRISPR targeting of Rbfox2*.*** Western blot shows lanes from the same gel, with staining for Gapdh and Rbfox2 in cells with infection with Cas9 and guide RNA (control) or with infection of Lenti_v2 containing Cas9 and a guide targeting Rbfox2.

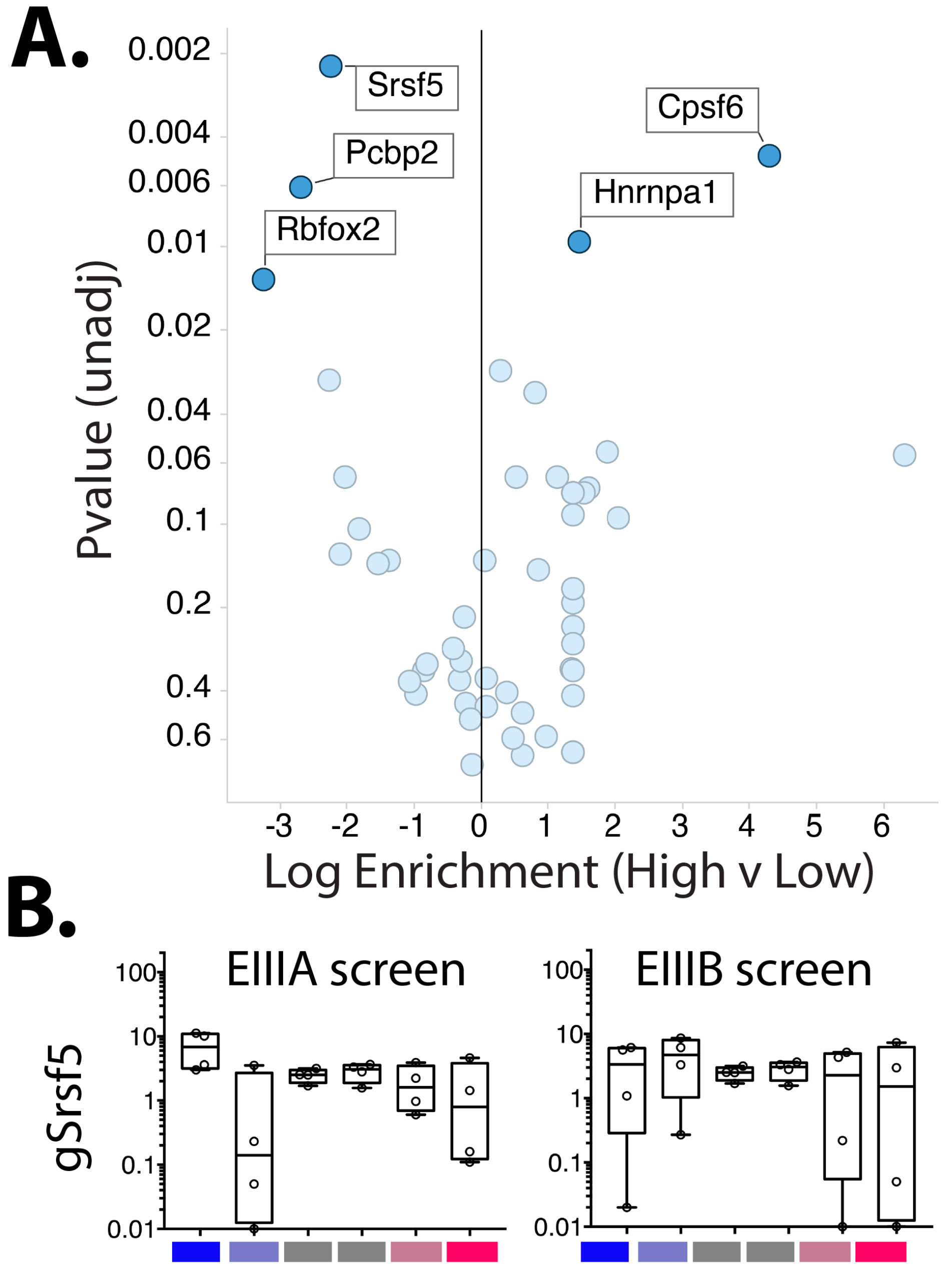


**SI Figure 5. Small scale screen for regulators of EIIIA.** (A) Volcano plot showing the enrichment of guides for the indicated genes among cells with high levels of EIIIA inclusion versus low levels of EIIIA inclusion. Log enrichment and p-value is from MaGeCK analysis. (B) Plots showing individual guides for Srsf5, and their relative enrichments in sorted cells with low, mid or high levels of EIIIA and EIIIB inclusion (blue to red, respectively).

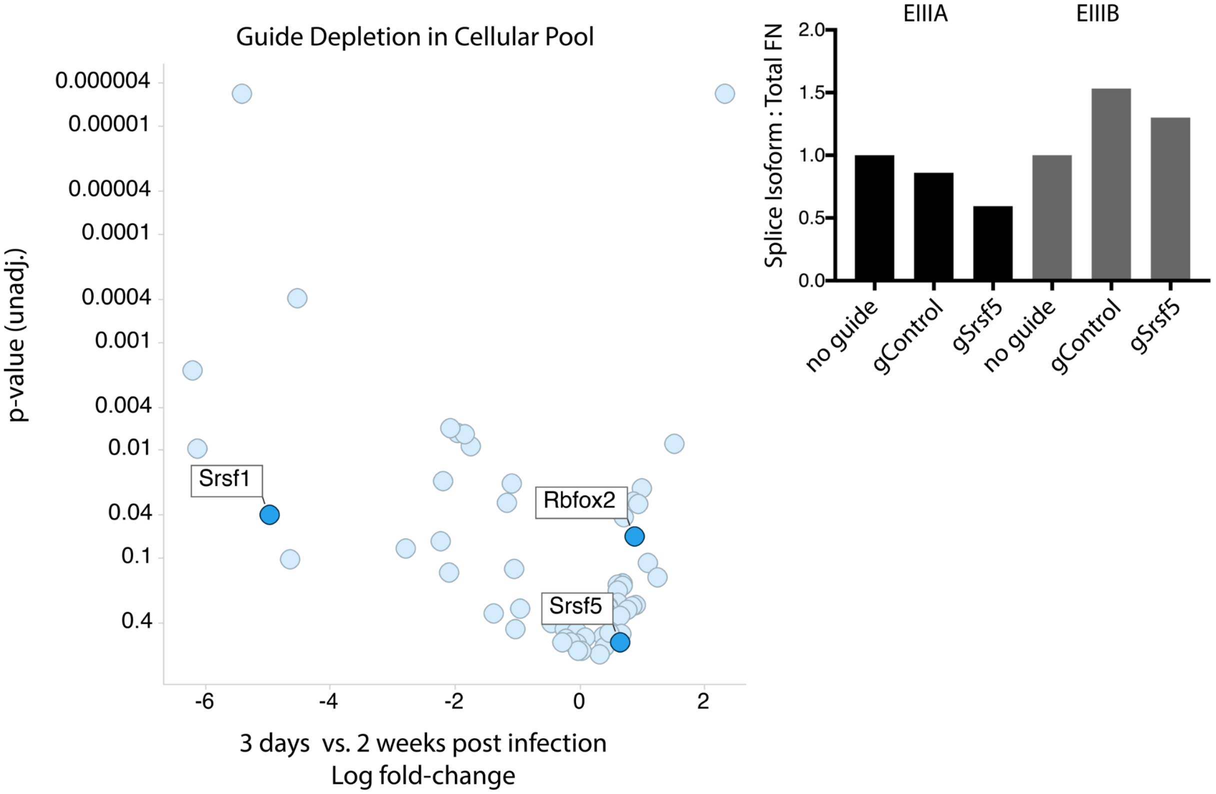


**SI Figure 6. Effects of deletion of Srsf1 and Srsf5 on EIIIA splicing.** (A) Volcano plot shows the change in guide representation in the cellular pool over time, and p-value by MaGeCK analysis. (B) qPCR analysis of the effect of targeting Srsf5 in cells on EIIIA inclusion level. The level of EIIIA specific PCR product relative to total FN is shown. Cells were not selected on the basis of deletion, so the population will contain some cells without Srsf5 deletion but with gSrsf5 and Cas9.

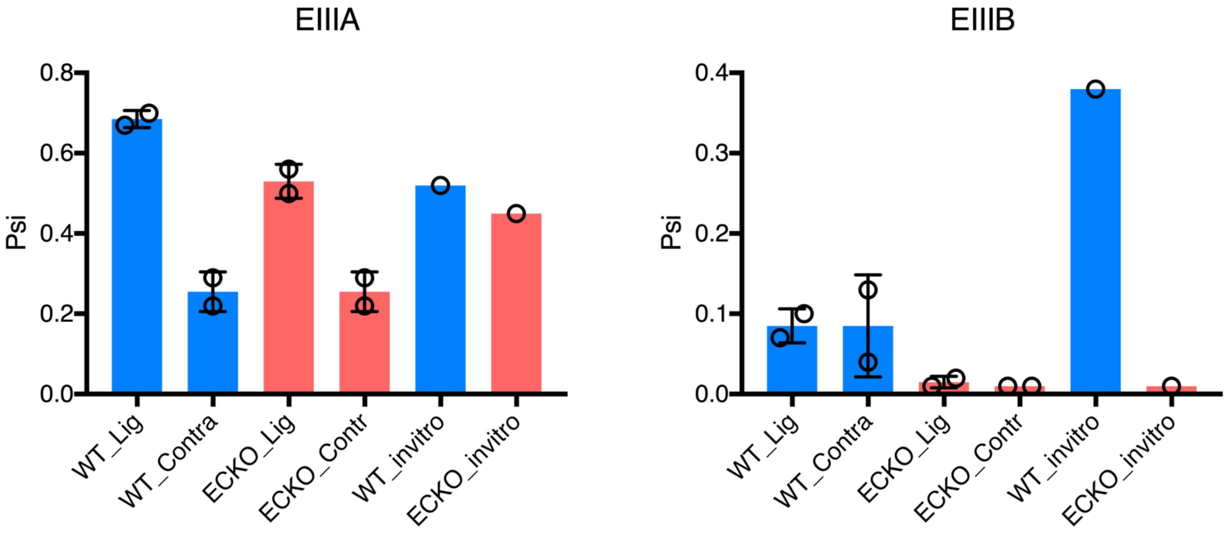


**SI Figure 7. Rbfox2 regulates EIIIA and EIIIB in aortic endothelial cells *in vivo* and *in vitro.*** Graphs show EIIIA and EIIIB inclusion in FN transcripts by MISO, *in vivo* or *in vitro*. Blue are wild-type controls (Rbfox2 ff) and red are Rbfox2 EC-KO mice (Cdh5(PAC)-CreERT2; Rbfox2 ff). *In vivo* data is from endothelial flush from carotid arteries exposed to low and disturbed flow for 7 days (WT_Lig and ECKO_Lig) or the contralateral arteries exposed to normal flow (WT_Contra and ECKO_Contra). *In vitro* cells were isolated from abdominal aorta and cultured *in vitro* under pro-inflammatory static conditions in the presence of serum for 7 days before sorting AcetylatedLDL+CD31+Icam2+ endothelial cells by flow cytometry.

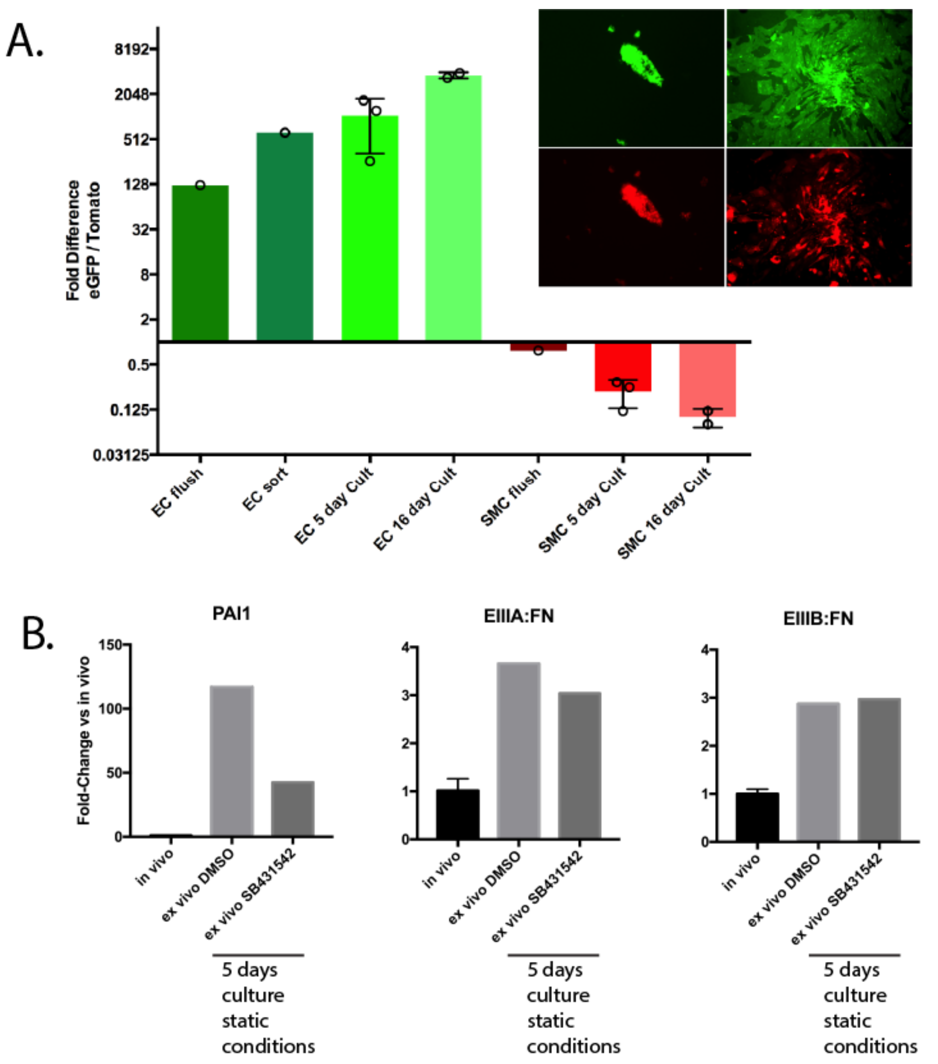


**SI Figure 8. Limited effect of TGFb receptor inhibition on EIIIA and EIIIB inclusion in aortic endothelial cells***.* Analysis of mRNA isolated from aortic endothelial cells in vivo, or after 5 days of culture under static conditions in vitro. (A) Cdh5(PAC)-CreERT2; mT/mG mice, were treated with tamoxifen to induce eGFP reporter expression in endothelial cells, and RNA was isolated by Trizol flush of the arterial vessel, or by sorting eGFP+ cells after in vitro culture. Image inset shows typical collagenase flush of intima from artery, which can be purified into endothelial (EC) or non-endothelial (mainly smooth muscle, SMC) by expression of eGFP. Graph shows the expression of eGFP (Cre-induced) vs tdTomato (no Cre) in RNA isolated from intimal artery flush, sorted endothelial cells (eGFP+) or sorted non-endothelial cells (eGFP neg). (B) Cells were cultured in the presence or absence of TGFb receptor (Alk5) inhibitor SB431542 or DMSO control for 5 days. PAI1 is a transcriptional target of TGFb signaling via Alk5. EIIIA:FN and EIIIB:FN refer to the ratio of FN transcripts containing the splice isoforms.

### SI Table 1. Sorted cell numbers used in each CRISPR screen

| Screen | Cell population | Number of Replicates | Cells/Replicate |
|---|---|---|---|
| Targeted (Fn-A) | Lowest 10% | 1 | 9482 |
| Targeted (Fn-A) | Low 10-20% | 1 | 10652 |
| Targeted (Fn-A) | Unsorted | 1 | ~1M |
| Targeted (Fn-A) | High 10-20% | 1 | 10814 |
| Targeted (Fn-A) | Highest 10% | 1 | 10414 |
| Targeted (Fn-B) | Lowest 10% | 1 | 10583 |
| Targeted (Fn-B) | Low 10-20% | 1 | 15815 |
| Targeted (Fn-B) | Unsorted | 1 | ~1M |
| Targeted (Fn-B) | High 10-20% | 1 | 10078 |
| Targeted (Fn-B) | Highest 10% | 1 | 10089 |
| Genome-wide (Fn-A) | Low 10% | 3 | ~267,000 |
| Genome-wide (Fn-A) | Mid 20% | 6 | ~775,000 |
| Genome-wide (Fn-A) | High 10% | 3 | ~254,000 |
| Genome-wide (Fn-B) | Low 10% | 3 | ~211,000 |
| Genome-wide (Fn-B) | Mid 20% | 6 | ~653,000 |
| Genome-wide (Fn-B) | High 10% | 3 | ~191,700 |
